# Simulating the Monty Hall problem in a DNA sequencing machine

**DOI:** 10.1101/502435

**Authors:** Noam Mamet, Gil Harari, Adva Zamir, Ido Bachelet

## Abstract

The Monty Hall problem is a decision problem with an answer that is surprisingly counter-intuitive yet provably correct. Here we simulate and prove this decision in a high-throughput DNA sequencing machine, using a simple encoding. All possible scenarios are represented by DNA oligonucleotides, and gameplay decisions are implemented by sequencing these oligonucleotides from specific positions, with a single run simulating more than 12,000,000 independent games. This work highlights high-throughput DNA sequencing as a new tool that could extend existing capabilities and enable new encoding schemes for problems in DNA computing.

---

A central question within the mathematical and computational contexts of DNA computing^1–10^ is the applicability of this field to real world problems. While DNA computing is currently unmatched by silicon-based computers in biological settings^11–17^, it is still not clear whether it could be adapted to solve very complex problems, e.g. NP-hard problems, at realistic scales, where conventional computers perform profoundly better. The limitations of DNA computing have been previously discussed^18^, but it is important to note that these limitations apply only for specific encoding schemes of the problem and methods of readout. The original scheme used by Adleman^1^ for a 7-vertex Hamiltonian Path problem, although innovative at the time and elegant as a proof of principle, was grossly non-scalable; however present day DNA synthesis and sequencing technologies could enable new schemes that address this issue at least in specific problems.

High-throughput DNA sequencing offers a unique setting to test diverse forms of encoding and readout that utilize the mechanics of the sequencing strategy being used^19^. For example, the Illumina technology sequences DNA by using sample strands as templates for *denovo* DNA synthesis^20^, which is performed on billions of unique sample strands in parallel. Each polymerization step is monitored in every strand by the sequencing machine. Thus, a typical sequencing run consists of hundreds of billions of individually-traced polymerization reactions. Moreover, older sequencers, such as the Illumina Genome Analyzer (GA)IIx, allow flexibility in designing recipes and introducing components to the chemistry and hardware of the machine, in turn enabling high-throughput screening of DNA properties such as protein binding^21^. Additional reactions, such as amplification reactions, restriction, ligation, editing, and re-sequencing, can be performed within the sequencing chamber, extending potential encoding schemes and providing readout that is highly resolved both spatially and temporally. This high level of parallelization is useful for many specific types of calculations.

Here, we used high-throughput DNA sequencing to simulate and numerically prove the correct answer to the Monty Hall problem^22^ (MHP). The essential layout of this problem is described in **Fig. 1A-C**. The participant, or subject, is presented with 3 closed doors. In the famous version of MHP, behind one of the doors is a new car, while each of the other two doors conceals a goat. The process then goes as follows:

1. The participant is asked to choose one of the doors, which remains closed **(Fig. 1A)**;
2. The presenter, who has complete knowledge of what is behind each door, opens a different door and reveals a goat **(Fig. 1B)**;
3. The participant should now decide whether to switch to the remaining closed door, or to stay with the initially-chosen one **(Fig. 1C)**;
4. The presenter opens the door finally chosen by the participant, and reveals what is behind it.

**Figure 1.**
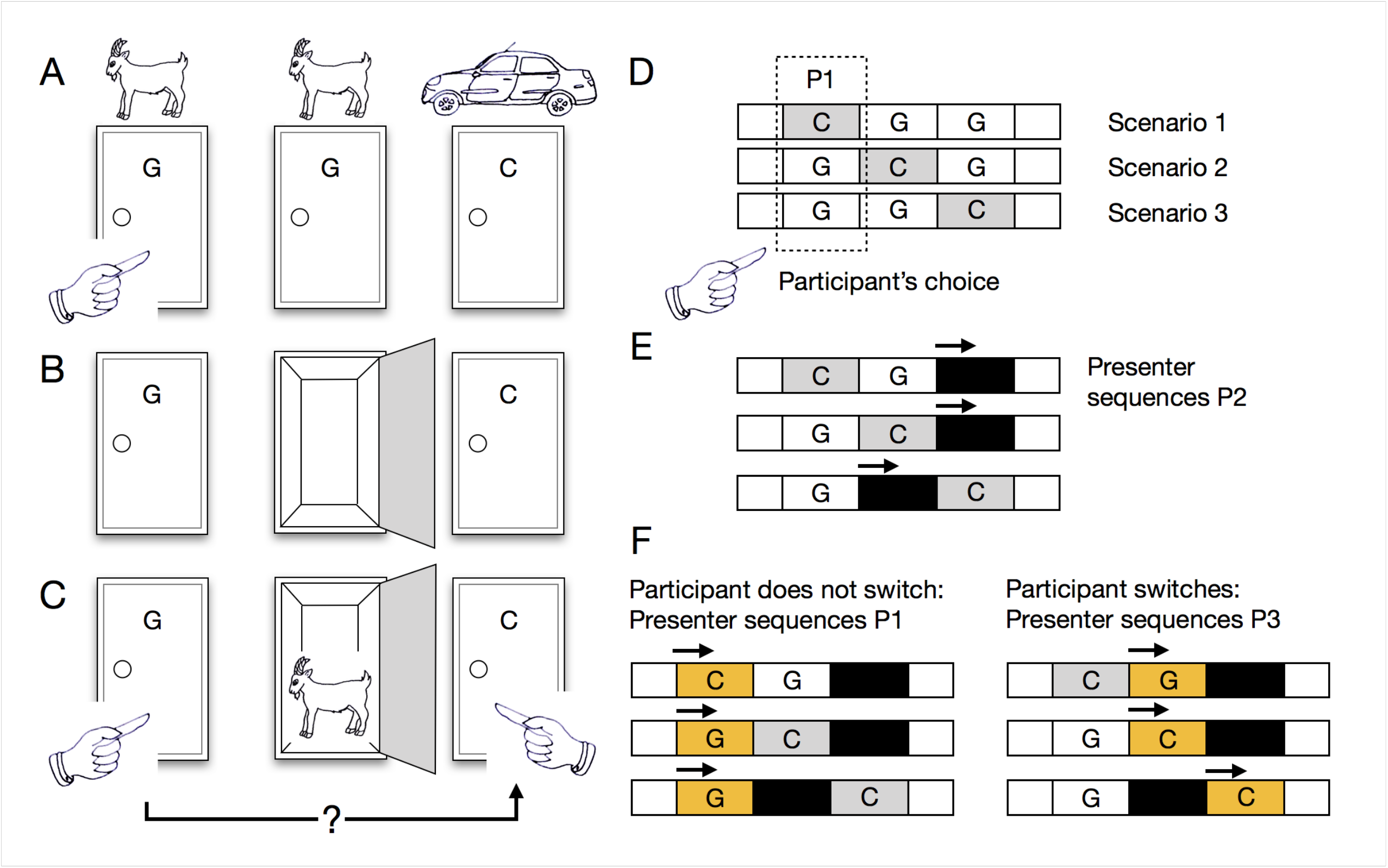
Layout of the Monty Hall problem (MHP) and its encoding in DNA molecules. **A-C**, a schematic of the MHP gameplay, showing a specific scenario (GGC). **A**, the participant is asked to choose one of the doors, which remains closed. **B**, the presenter opens a door thereby intentionally revealing a goat. **C**, the core choice is then made by the participant whether to switch to the other closed door or not to switch. **D**, encoding of the MHP in an ensemble of three DNA oligonucleotides, which represent all possible scenarios of the game (CGG, GCG, GGC). The primer P1 represents the participant’s first choice across the ensemble. **E**, the presenter’s action of opening a door which reveals a goat is encoded by sequencing the entire ensbemble from the primer P2. **F**, the decision to switch is encoded by sequencing the ensemble again from either the primer P1 (first choice) or the primer P3 (the choice to switch). Results of the decision are revealed as sequences of the identifier barcodes.

The core of this decision problem is contained in step 3, namely, should the participant switch to the remaining (unchosen and unopened) third door, or to remain with the door that was chosen originally. It can be proven mathematically that the correct decision is to switch to the third door, which leads to the best chance of winning the car. In step 1, the probability of the car being behind the chosen door is ⅓, and of it being behind either of the other two doors is collectively ⅔; however, step 2 shows that one of these two doors is worthless, effectively shifting the entire probability of ⅔ to the remaining door. Switching doors therefore doubles the probability for winning the car from ⅓ to ⅔.

Despite the fact it can be proven, the correct answer is surprisingly counter-intuitive. The decision between only two doors strongly suggests that each choice should be assigned a probability of ½, in which case switching and not switching have equal value. One approach to demonstrate the paradoxical nature of this intuition, and subsequently prove the correct answer, is to run many iterations of MHP and show that the decision to switch leads to reward in ⅔ of the cases. This simulation was the one performed in this work, using the encoding scheme described below.

In order to simulate all the possible outcomes of MHP, we designed DNA oligonucleotides that contain 3 regions, each region representing a car (C) or a goat (G). The oligonucleotide ensemble contained every possible configuration of the 3 doors (CGG, GCG, and GGC) **(Fig. 1D)** in equal molar ratios. Each region in any of the configurations is built from two parts: an access primer (P1, P2, P3) and an 8-nt identifier barcode (B1 for C, B2 for G).

In order to capture the unique structure of MHP, the access primers in this implementation are defined in terms of gameplay decisions and not physical position, i.e. they do not represent a specific “door”. The first primer (P1) represents the participant’s first choice, which may be a car (probability of ⅓) or either goat 1 or 2 (collective probability of ⅔). The second primer (P2) represents the elimination of a goat by the presenter **(Fig. 1E)**, and the third primer (P3) represents the participant’s decision to switch **(Fig. 1F)**.

There are three different scenarios in MHP **(Table 1)**. The participant may randomly choose the car, goat number 1, or goat number 2. In the first case, if the participant chooses the car, the presenter reveals a goat leaving the other goat for the participant in case a decision to switch is made. In the other two scenarios, a goat has been chosen and the presenter will reveal the second goat, leaving only the car in case the participant decides to switch. In our implementation of MHP, each oligonucleotide represents one of the three scenarios. The specific design used here is as follows, although the particular order of regions along each oligonucleotide is irrelevant:

1. Scenario 1 = [C][G][G] = [P1B1][P3B2][P2B2]
2. Scenario 2 = [G][C][G] = [P1B2][P3B1][P2B2]
3. Scenario 3 = [G][G][C] = [P1B2][P2B2][P3B1]

**Table 1.**
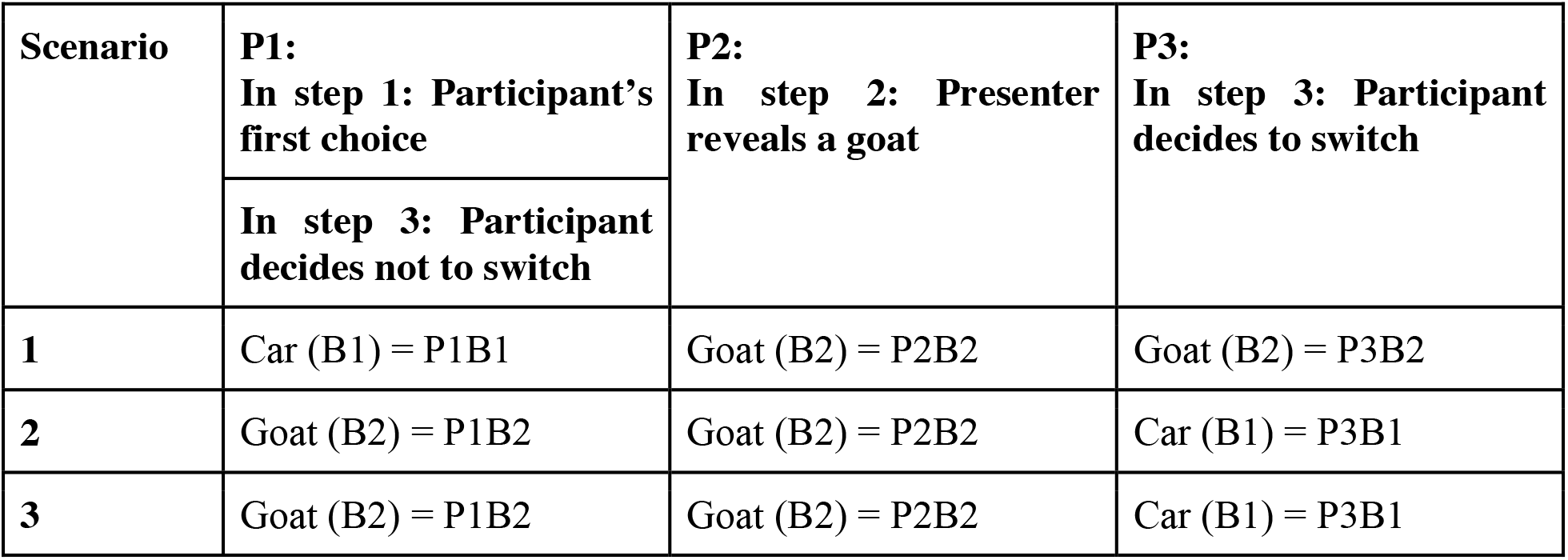
Detailed structure of the MHP ensemble, capturing all possible scenarios of the game.

| Scenario | **P1:**<br>In step 1: Participant's first choice | **P2:**<br>In step 2: Presenter reveals a goat | **P3:**<br>In step 3: Participant decides to switch |
| --- | --- | --- | --- |
|  | In step 3: Participant decides not to switch |  |  |
| **1** | Car (B1) = P1B1 | Goat (B2) = P2B2 | Goat (B2) = P3B2 |
| **2** | Goat (B2) = P1B2 | Goat (B2) = P2B2 | Car (B1) = P3B1 |
| **3** | Goat (B2) = P1B2 | Goat (B2) = P2B2 | Car (B1) = P3B1 |

This simple encoding captures the precise structure of MHP.

The MHP ensemble was sequenced in an Illumina NextSeq 500 instrument in paired-end mode. In the first read, the full length of the ensemble was sequenced to ensure that all scenarios are equally represented **(Fig. 2A)**; the second read was sequenced from a specific access primer (P1, P2, or P3) based on gameplay decision.

**Figure 2.**
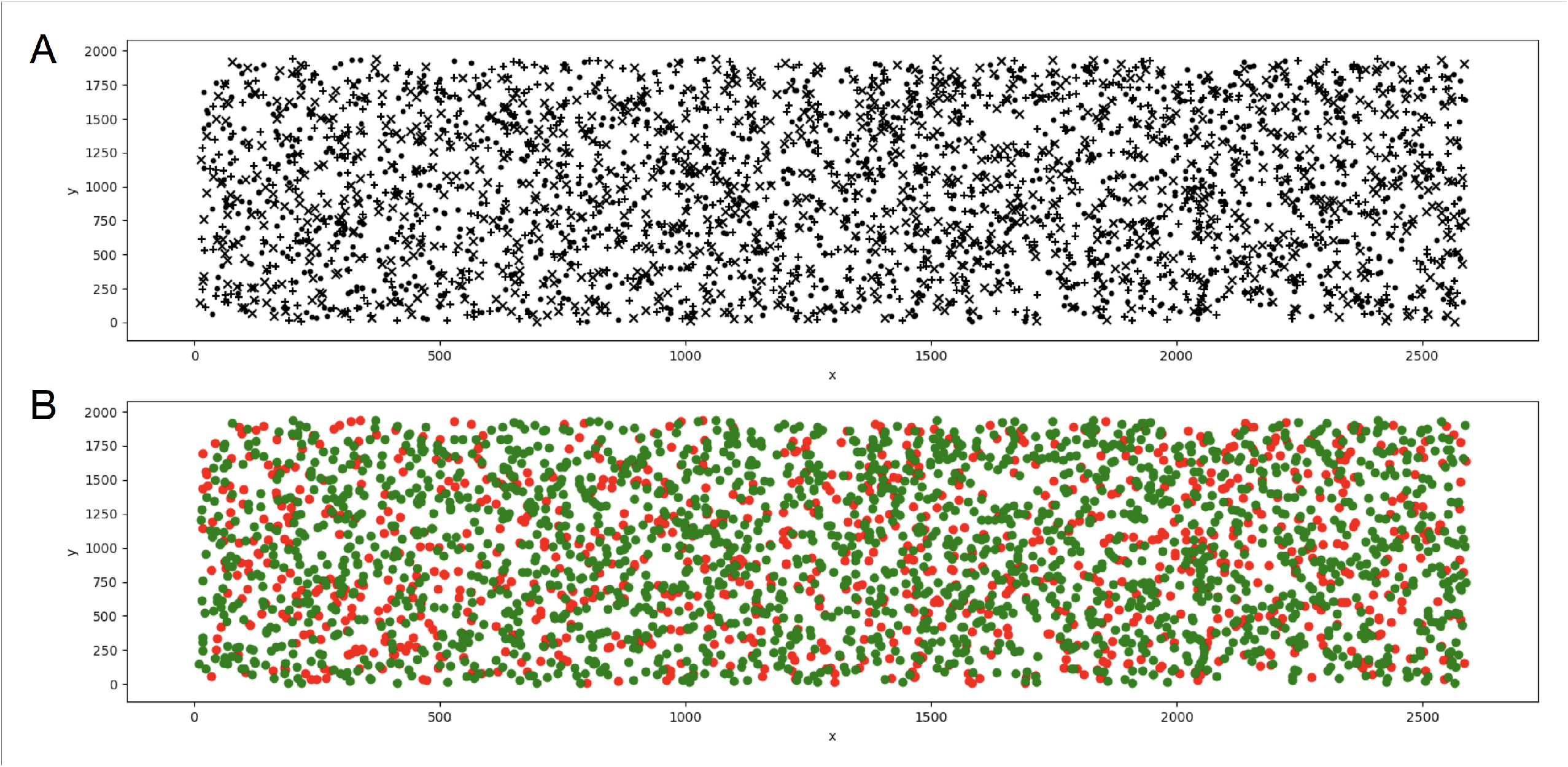
Experimental results of the MHP simulation. **A**, a representative field from a flow-cell tile displaying 3,000 clusters in 2D space; only every fifth cluster in the field is displayed for purposes of clarity. Each of the three symbols (x, +, dot) represents a scenario from the ensemble. X and Y units are pixels, using the native Illumina NextSeq image resolution of 0.8 microns/pixel. Image dimensions are 2592 × 1944 pixels. **B**, results of sequencing the ensemble following the participant’s decision to switch doors. Sequencing took place from primer P3 instead of P1 (the first choice in the game), and barcode sequences were analyzed and represented by colors (green, car barcode; red, goat barcode). The run resulted in 8,227,528 “cars”, which is 66.7% of the total number of 12,341,292 games that were simulated.

Following the first read, the experimental procedure progressed as follows:

1. The participant’s chose a “door”, with a probability of ⅓ to conceal a car and ⅔ to conceal a goat; this choice, which is always the same in MHP regardless of scenario, was then defined by P1, but was not sequenced at this stage.
2. The presenter sequenced the ensemble from access primer P2, revealing a goat across the entire ensemble.
3. The participant chose to switch: the presenter sequenced the ensemble from access primer P3.

Note: had the participant chosen not to switch, presenter would have sequenced from access primer P1.

This run simulated 12,341,292 games, and the decision to switch resulted in a car in 8,227,528 games, which is 66.7% of the total well-sequenced reads **(Fig. 2B)**.

The purpose of this work was to highlight high-throughput DNA sequencing as a setup to probe new possibilities in the encoding of problems that rely on parallel computing in order to be solved. Moreover, it shows one possible approach in the representation of such problems. The technology employed here is routinely used to sequence billions of reads with more than 100 bases each. Even as a readout mechanism alone, this already significantly expands the capabilities of early schemes such as Adleman’s. In this scheme^1^, in which each vertex and each edge is represented by a 20-mer oligonucleotide, a graph with *k* vertices would require a total of 20*k*(log *k*)^*k*^ base pairs of DNA, reaching an unrealistic scale rapidly beyond the initial graph of 7 vertices. In contrast, high-throughput sequencing could theoretically support representations of graphs with 40–50 vertices using Adleman’s original encoding of the problem. Although this is still very far from the current capabilities of conventional computers, it demonstrates that what was considered once completely impractical can now potentially be achieved by an existing, off-the-shelf technology, and by improved encoding forms.

It would be interesting to see such new forms of computational problem encoding in DNA, that utilize the entire range of capabilities e.g. high-throughput sequencing, DNA editing, strand-displacement networks, etc., which became available only recently. For example, it could be feasible to develop a scheme for the division of large and complex problems into billions of small tasks, their distributed execution within the sequencing machine, and later their reintegration into a solution to the original problem, inside the sequencer. Benenson et al^8^ describe a finite automaton made of DNA molecules. This machine could be programmed to parform calculations. In their work, 10^12^ processes were run in a single tube, carrying out the same calculation. Implemented in a sequencing machine, 10^10^ automata could be programmed to perform 10^10^ unique calculations, each monitored independently.

Although it is still early to see where and to what extent would high-throughput DNA sequencing impact the question raised at the beginning of this article, it is not unlikely that further advances in DNA sequencing, which for the past decade significantly outpaced Moore’s law^23^, would lead to new insights into how can the current gaps be bridged.

## Methods summary

Oligonucleotides were ordered from Integrated DNA Technologies (IDT). Access primers were either taken directly from Nexterra and TruSeq sequencing primer collections, or custom designed, with T_m_ of ∼ 65 □. The barcodes were taken from the Nexterra barcode library. To compensate for the low complexity of the ensemble, an additional random 5-nt adapter was inserted at the beginning of each oligonucleotide, to allow the instrument to properly generate the cluster map during the first read. The concentration of each oligonucleotide was measured by Qubit and qPCR before mixing to obtain the ensemble. Sequencing library was prepared according to standard protocols. The ensemble was sequenced in an Illumina NextSeq 500 instrument in paired-end mode.

## Acknowledgements

The authors wish to thank Dana Bachelet for assistance with figures, and to the team at Augmanity for valuable discussions and assistance.

## Author contributions

N.M. and I.B. designed research and experiments and wrote the manuscript. N.M., G.H., and A.Z. performed experiments. N.M., G.H., and I.B. analyzed the data.

## Additional information

The authors declare that they have no competing interests.

## Supplementary Information

The complete ensemble is as follows:

Car in position 1:

AATGATACGGCGACCACCGAGATCTACACACACTCTTTCCCTACACGACGCTCTTCCGATCTNNNNNCTAGTACGGTTGCGTCACACTGAACATCCTTCTCTTGTGTGCCTAGTACGCTGTCTCTTATACACATCTCCGAGCCCACGAGACTCGCCTTACTGTCTCTTATACACATCTGACGCTGCCGACGAATCTCGTATGCCGTCTTCTGCTTG

Car in position 2:

AATGATACGGCGACCACCGAGATCTACACACACTCTTTCCCTACACGACGCTCTTCCGATCTNNNNNCTAGTACGGTTGCGTCACACTGAACATCCTTCTCTTGTGTGCTCGCCTTACTGTCTCTTATACACATCTCCGAGCCCACGAGACCTAGTACGCTGTCTCTTATACACATCTGACGCTGCCGACGAATCTCGTATGCCGTCTTCTGCTTG

Car in position 3:

AATGATACGGCGACCACCGAGATCTACACACACTCTTTCCCTACACGACGCTCTTCCGATCTNNNNNTCGCCTTACTGTCTCTTATACACATCTCCGAGCCCACGAGACCTAGTACGGTTGCGTCACACTGAACATCCTTCTCTTGTGTGCCTAGTACGCTGTCTCTTATACACATCTGACGCTGCCGACGAATCTCGTATGCCGTCTTCTGCTTG

Color coding:

Nexterra p5 adapter

TrueSeq p5 sequencing primer

Random adapter

“Presenter exposes a goat” sequencing primer (custom primer)

“Participant chooses to switch” sequencing primer (Nexterra p7 sequencing primer)

“Participant chooses not to switch” sequencing primer (Nexterra p5 sequencing primer)

Goat barcode

Car barcode

p7 complementary adapter

The ensemble was constructed as follows: each gameplay oligo was divided into two parts which were ordered separately (table 1). The two parts were hybridized as follows:

95°, 3min,
95°, 30 sec,
70.4°, 30 sec
72°, 30 sec,
6 Cycles

**Table S1:**
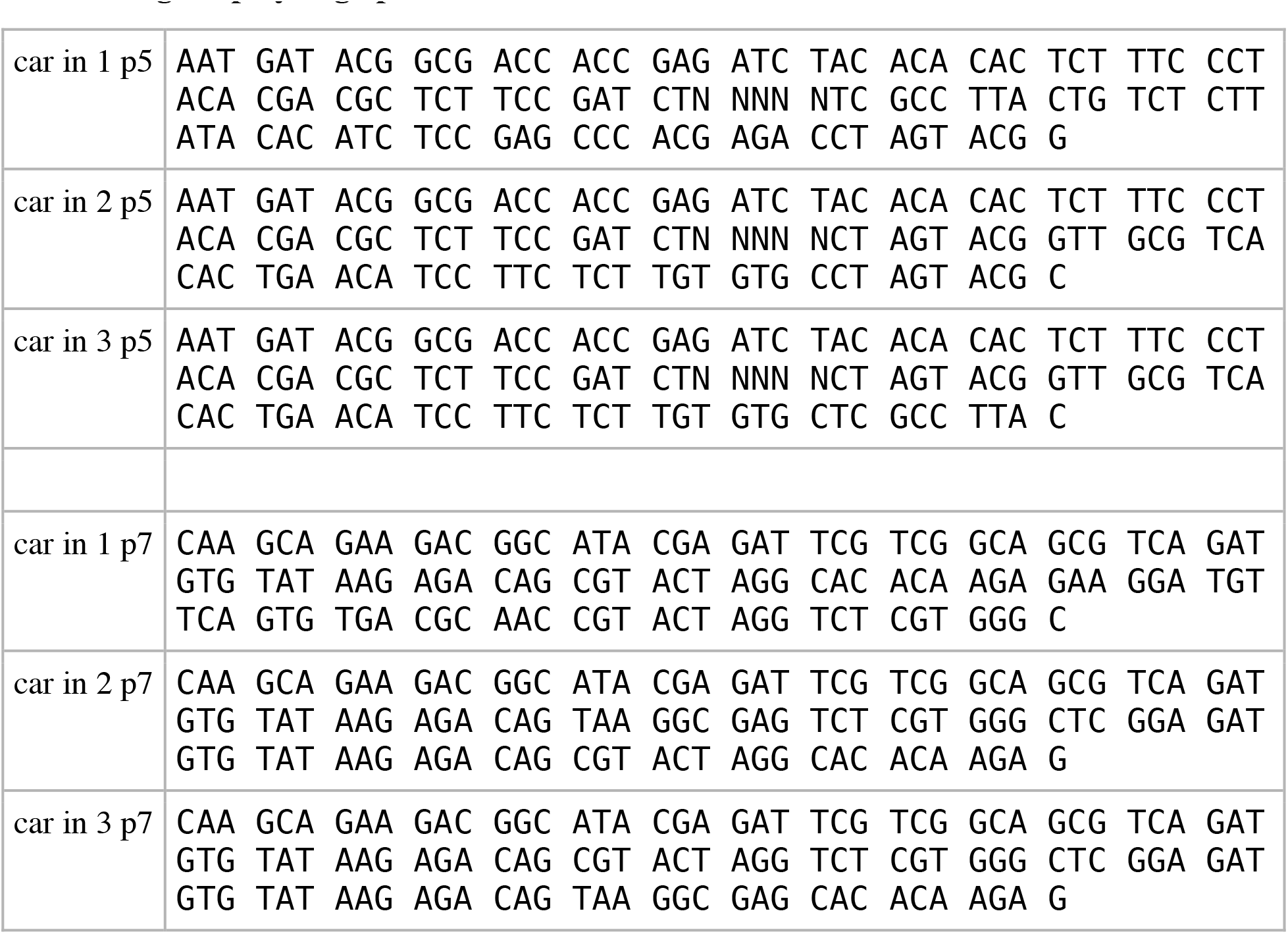
gameplay oligo parts.

**Table S2:**
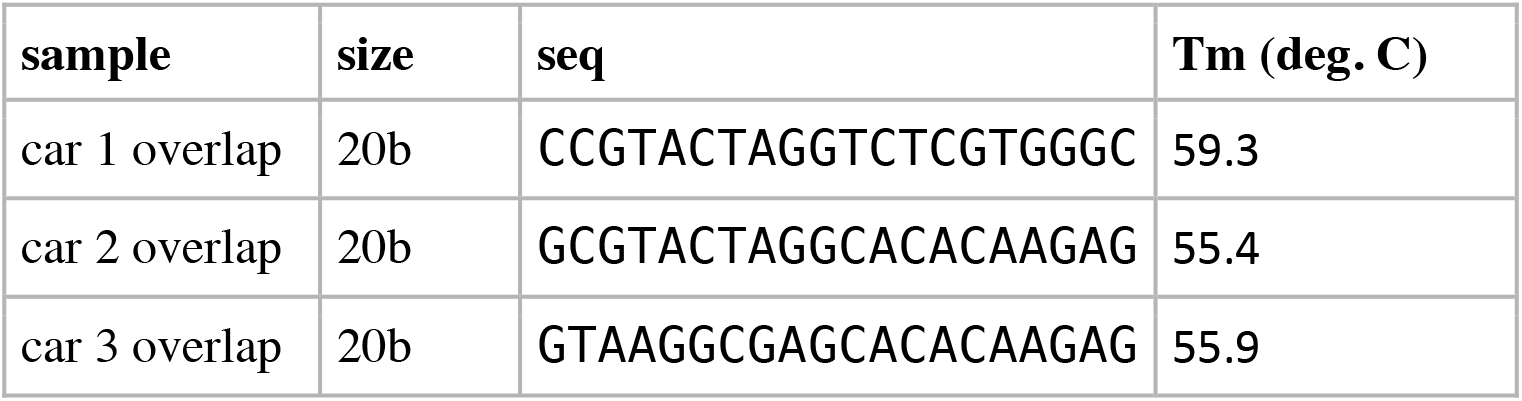
overlap in each pair.

Samples were measured in Qubit to verify concentration and purity. 10 independent mixtures were made, and measurements were carried out to verify mixing accuracy.

**Table S3:**
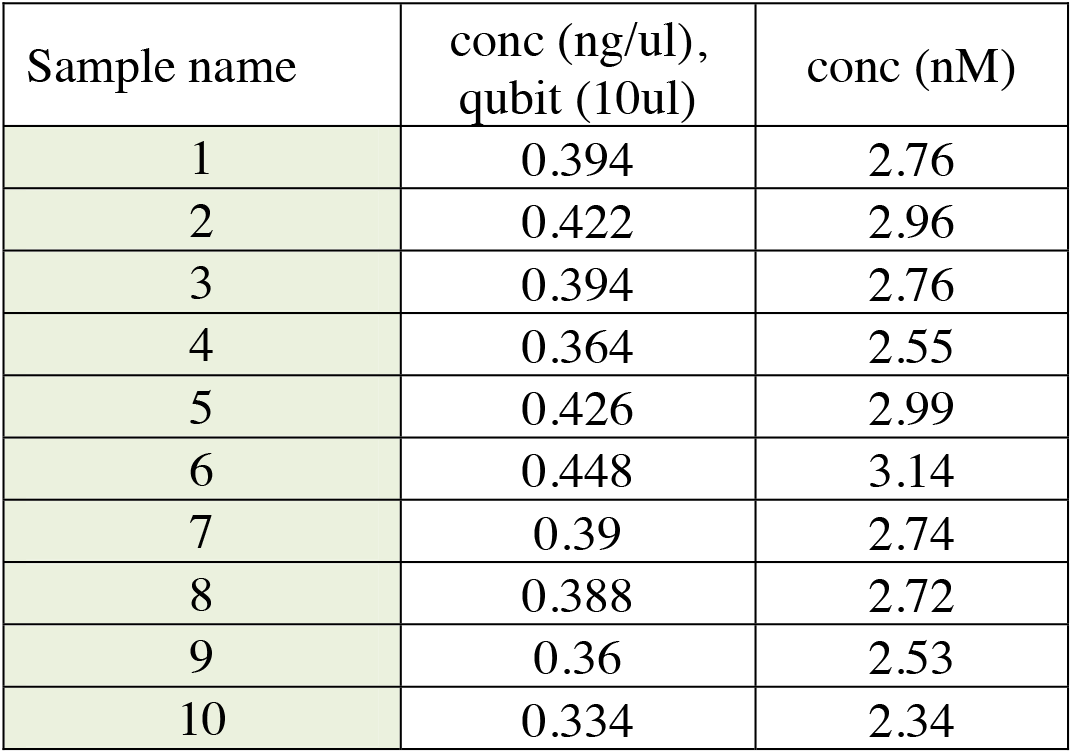
quality analysis.

1. qPCR was carried out on the 10 independent mixtures. Primers used were as follows:

For AATGATACGGCGACCAC
Rev CAAGCAGAAGACGGCATA

**Fig. S1:**
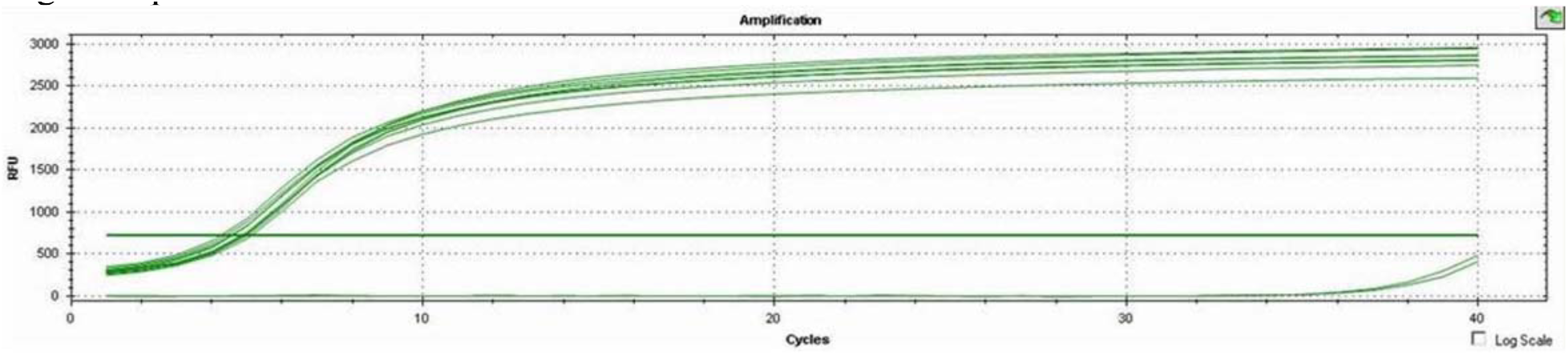
qPCR results.

