## Supplementary information for "Simulating the Monty Hall problem in a DNA sequencing machine"

The complete ensemble is as follows:

Car in position 1:

AATGATACGGCGACCACCGAGATCTACACACACTCTTTCCCTACACGACGCTCTTCCGATCTNN
NNNCTAGTACGGTTGCGTCACACTGAACATCCTTCTCTTGTGTGCCTAGTACGCTGTCTCTTAT
ACACATCTCCGAGCCCACGAGACTCGCCTTACTGTCTCTTATACACATCTGACGCTGCCGACGA
ATCTCGTATGCCGTCTTCTGCTTG

Car in position 2:

AATGATACGGCGACCACCGAGATCTACACACACTCTTTCCCTACACGACGCTCTTCCGATCTNN
NNNCTAGTACGGTTGCGTCACACTGAACATCCTTCTCTTGTGTGCTCGCCTTACTGTCTCTTAT
ACACATCTCCGAGCCCACGAGACCTAGTACGCTGTCTCTTATACACATCTGACGCTGCCGACGA
ATCTCGTATGCCGTCTTCTGCTTG

Car in position 3:

AATGATACGGCGACCACCGAGATCTACACACACTCTTTCCCTACACGACGCTCTTCCGATCTNN
NNNTCGCCTTACTGTCTCTTATACACATCTCCGAGCCCACGAGACCTAGTACGGTTGCGTCACA
CTGAACATCCTTCTCTTGTGTGCCTAGTACGCTGTCTCTTATACACATCTGACGCTGCCGACGA
ATCTCGTATGCCGTCTTCTGCTTG

Color coding:
Nexterra p5 adapter
TrueSeq p5 sequencing primer
Random adapter
“Presenter exposes a goat” sequencing primer (custom primer)
“Participant chooses to switch” sequencing primer (Nexterra p7 sequencing primer)
“Participant chooses not to switch” sequencing primer (Nexterra p5 sequencing primer)
Goat barcode
Car barcode
p7 complementary adapter

The ensemble was constructed as follows: each gameplay oligo was divided into two parts which were ordered separately (table 1). The two parts were hybridized as follows:

95°, 3min,
95°, 30 sec,
70.4°, 30 sec
72°, 30 sec,
6 Cycles

**Table S1: gameplay oligo parts**

| | |
|---|---|
| car in 1 p5 | AAT GAT ACG GCG ACC ACC GAG ATC TAC ACA CAC TCT TTC CCT ACA CGA CGC TCT TCC GAT CTN NNN NTC GCC TTA CTG TCT CTT ATA CAC ATC TCC GAG CCC ACG AGA CCT AGT ACG G |
| car in 2 p5 | AAT GAT ACG GCG ACC ACC GAG ATC TAC ACA CAC TCT TTC CCT ACA CGA CGC TCT TCC GAT CTN NNN NCT AGT ACG GTT GCG TCA CAC TGA ACA TCC TTC TCT TGT GTG CCT AGT ACG C |
| car in 3 p5 | AAT GAT ACG GCG ACC ACC GAG ATC TAC ACA CAC TCT TTC CCT ACA CGA CGC TCT TCC GAT CTN NNN NCT AGT ACG GTT GCG TCA CAC TGA ACA TCC TTC TCT TGT GTG CTC GCC TTA C |
| car in 1 p7 | CAA GCA GAA GAC GGC ATA CGA GAT TCG TCG GCA GCG TCA GAT GTG TAT AAG AGA CAG CGT ACT AGG CAC ACA AGA GAA GGA TGT TCA GTG TGA CGC AAC CGT ACT AGG TCT CGT GGG C |
| car in 2 p7 | CAA GCA GAA GAC GGC ATA CGA GAT TCG TCG GCA GCG TCA GAT GTG TAT AAG AGA CAG TAA GGC GAG TCT CGT GGG CTC GGA GAT GTG TAT AAG AGA CAG CGT ACT AGG CAC ACA AGA G |
| car in 3 p7 | CAA GCA GAA GAC GGC ATA CGA GAT TCG TCG GCA GCG TCA GAT GTG TAT AAG AGA CAG CGT ACT AGG TCT CGT GGG CTC GGA GAT GTG TAT AAG AGA CAG TAA GGC GAG CAC ACA AGA G |

**Table S2: overlap in each pair**

| **sample** | **size** | **seq** | **Tm (deg. C)** |
|---|---|---|---|
| car 1 overlap | 20b | CCGTACTAGGTCTCGTGGGC | 59.3 |
| car 2 overlap | 20b | GCGTACTAGGCACACAAGAG | 55.4 |
| car 3 overlap | 20b | GTAAGGCGAGCACACAAGAG | 55.9 |

Samples were measured in Qubit to verify concentration and purity. 10 independent mixtures were made, and measurements were carried out to verify mixing accuracy.

**Table S3: quality analysis**

| Sample name | conc (ng/ul), qubit (10ul) | conc (nM) |
|---|---|---|
| 1 | 0.394 | 2.76 |
| 2 | 0.422 | 2.96 |
| 3 | 0.394 | 2.76 |
| 4 | 0.364 | 2.55 |
| 5 | 0.426 | 2.99 |
| 6 | 0.448 | 3.14 |
| 7 | 0.39 | 2.74 |
| 8 | 0.388 | 2.72 |
| 9 | 0.36 | 2.53 |
| 10 | 0.334 | 2.34 |

1. qPCR was carried out on the 10 independent mixtures. Primers used were as follows:

For AATGATACGGCGACCAC
Rev CAAGCAGAAGACGGCATA

Fig. S1: qPCR results

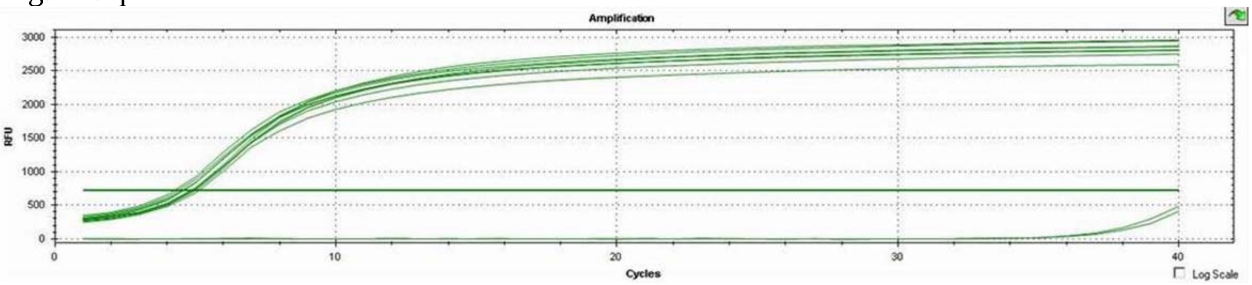
